# Harnessing extracellular vesicles for efficient siRNA delivery *in vitro* and *in vivo*

**DOI:** 10.1101/2023.08.26.554924

**Authors:** Tao Qiu, Yanhua Zhai, Yuan Yi, Xiaoyu Wang, Rui Hu, Shuiqin Niu, Chengcheng Wang, Chuang Cui, Xinjun He, Ke Xu

**Affiliations:** Vesicure Therapeutics Co. Ltd., Biobay 3B, Suzhou Industrial Park, Suzhou, 215000, China

## Abstract

To fulfill the potential of small interference RNA (siRNA) therapeutics, diverse and efficient delivery platforms for siRNA are urgently needed. Extracellular vesicles (EVs) are endogenous, cell secreted nano-vehicles which have been explored for functional siRNA delivery, yet impurities in EVs preparation and lack of efficient siRNA loading method have been limiting its further development into clinical applications. In this work, we have achieved high quality EVs production and an siRNA loading system via EVs. The prepared EV-siRNA system was able to deliver siRNA cargo to model cell lines to silence the expression of targeted genes with high potency. Furthermore, EV-siRNA administration in a tumor xenograft model significantly inhibited tumor growth. Therefore, this study establishes an innovative and efficient EV-siRNA platform for siRNA delivery *in vitro* and *in vivo*.

## Introduction

Small interference RNA (siRNA) therapeutics has been rapidly emerged as an innovative modality of medicines (Hu et al. 2020; Friedrich and Aigner 2022). siRNA functional delivery involves transporting small RNA molecules into cells to silence specific genes and regulate protein expression. It holds promise for targeting various diseases by targeting disease-causing genes. This technology has revolutionized gene therapy and research, however, it comes with certain challenges, limiting the clinical translation of siRNA therapy: siRNAs have difficulty entering cells by themselves due to their size and negative charge, necessitating efficient delivery methods. Currently, GalNAc (N-acetylgalactosamine) and LNPs (Lipid Nanoparticles) are two prominent approaches for siRNA delivery, with five drugs already approved by FDA, and a few under clinical trials evaluation (Friedrich and Aigner 2022). GalNAc technology is highly effective for liver-targeted siRNA delivery with lower doses and reduced off-target effects, but it’s limited to liver-related applications (Springer and Dowdy 2018). LNPs, on the other hand, offer higher delivering capacities, but their efficacy can be influenced by formulation and potential immunogenicity (Yonezawa, Koide, and Asai 2020). Therefore, innovative siRNA delivery platforms with extra-hepatic targeting ability and lower immunogenicity are urgently needed.

Extracellular vesicles (EVs) are nanometer sized vesicles (40-200 nm) secreted by most cell types and function as important mediators of intercellular communication (Kowal, Tkach, and Thery 2014). During biogenesis, EVs naturally package proteins, peptides, nucleic acids, lipids and other biomolecules, thus are already being investigated as delivery platform of various therapeutics (Vader et al. 2016). As drug delivery vehicles, EVs offer many advantages such as low immunogenicity and the potential of extra-hepatic targeting ability either by naturally occurring or artificial modifications (Zeng et al. 2023). Therefore, we are intrigued to explore the potential of EVs for siRNA delivery.

To date, EVs have already been demonstrated of siRNA delivery ability in several studies (Alvarez-Erviti et al. 2011; Kumar et al. 2015; Kamerkar et al. 2017; Amiri et al. 2022). Nonetheless, in majority of those work, we note critical shortcomings that may prevent the future translation from laboratory research into clinical applications. The first one is that EVs are mostly isolated through ultracentrifugation, due to the limited availability of cell culture supernatants as starting material especially for source cells such as immune cells and mesenchymal stem cells (MSC) under laboratory scale. Therefore, substantial impurities such as cellular debris and aggregated protein particles are likely present in the EVs fraction acquired for testing. Secondly, electroporation is currently the main method for siRNA loading into EVs, which transiently create holes in EV membranes, thereby allowing free siRNA diffusing into EV lumen (El-Andaloussi et al. 2012). On one hand, electroporation has been proved to cause siRNA aggregates and precipitates on EV surface, not only decreasing the loading efficiency but also further compromising EVs quality (Kooijmans et al. 2013). On the other hand, electroporation is not an easily scalable method, thus limiting the large-scale batch production.

In this work, we set out to optimize the production, purification and siRNA loading of EVs, with high quality and in a scalable and consistent way. We further demonstrated the efficient EV-siRNA delivery in both *in vitro* and *in vivo* models.

## Results

### Production, purification and characterization of HEK293F-derived extracellular vesicles

HEK293F cells adapted for suspension culture are chosen as the source of EVs production, since it can be cultured to high density, providing sufficient starting materials for density gradient based purification of EVs (**Figure 1A**). The purified EVs were enriched for common EV markers such as CD9, CD63, CD81 and TSG101 (**Figure 1B**). Nanoparticle tracking analysis (NTA) performed for multiple EVs batches revealed that isolated EVs had average size around 126 nm (**Figure 1C**). Importantly, purified EVs under transmission electron microscopy (TEM) presented minimal contaminants, suggesting of extreme high purity and quality (**Figure 1D**). In summary, we are able to consistently acquire large amount of high quality HEK293-EVs.

**Figure 1.**
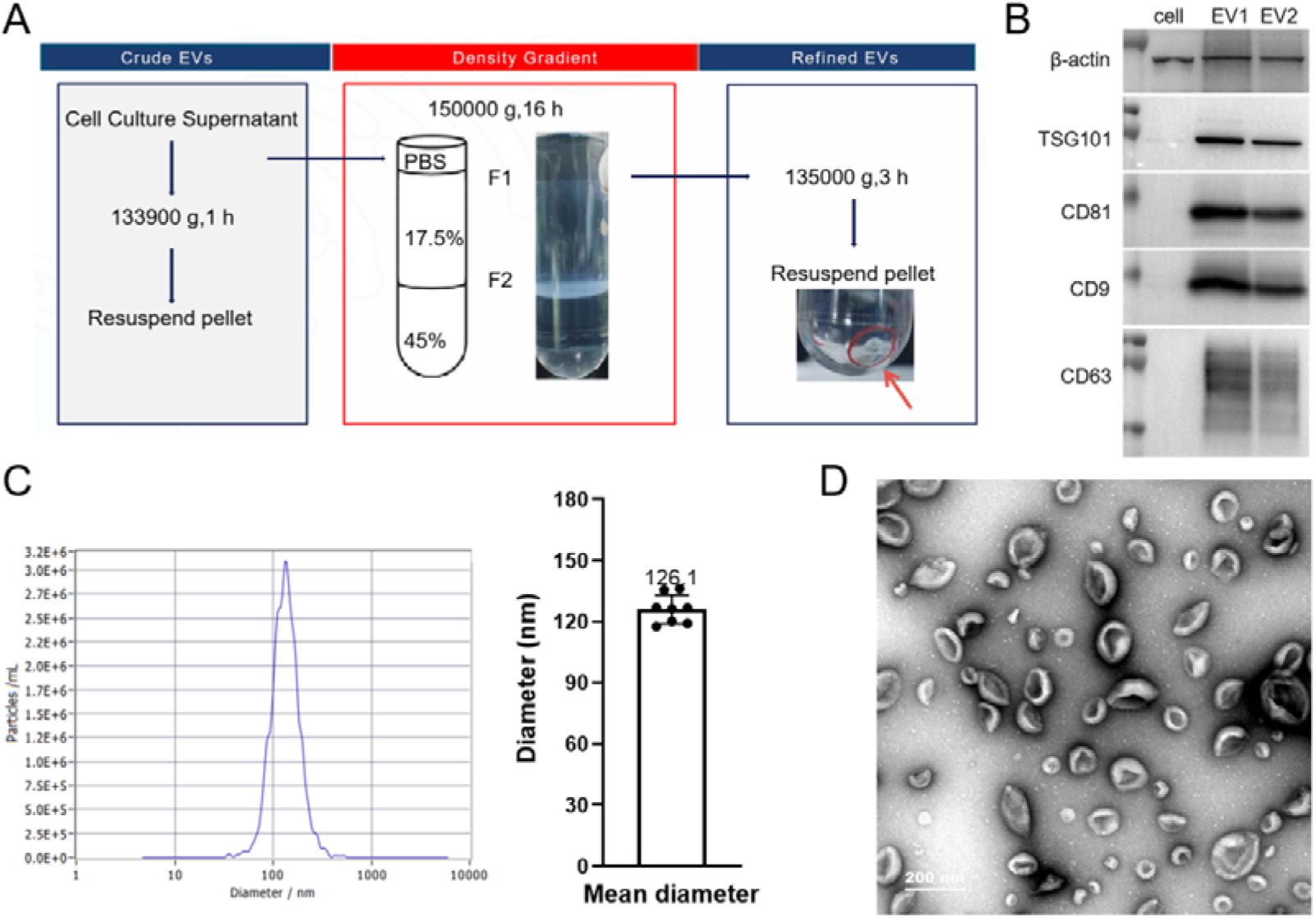
Production, purification and characterization of HEK293F-derived extracellular vesicles. (A) Schematic of density gradient based purification of HEK293F-derived EVs. (B) Whole cell lysates and EVs lysates from two batch of EVs preparation (EV1 and EV2) were analyzed for enrichment of common EVs markers including CD9, CD63, CD81 and TSG101. (C) Representative nanoparticle tracking analysis (NTA) analysis of size distribution and concentration of purified EVs. The mean diameter from eight different batches of EVs were presented. (D) Representative transmission electron microscopy image of purified EVs. Scale bar: 200 nm.

### Extracellular vesicles loading of siRNA and characterization

We utilized a home-developed loading system, dilEVry™, for high density siRNA loading into EV lumen. NTA analysis showed that following siRNA loading, EVs had a slightly increased mean size around 136 nm (**Figure 2A**). Next, EV-siRNA were tested for siRNA delivery ability. Efficient cellular uptake of EV-siRNA was observed in various cell types including a human lung cancer tumor cell line (H358), a human endothelial primary cell type (HUVEC) and a mouse macrophage cell line (RAW264.7) (**Figure 2B**). To evaluate the potential toxicity of EV-siRNA, CCK-8 assay was performed for cells at 72 hours post EV-siR-Scramble transfection, which showed that EV-siRNA were non-toxic and safe to the cells (**Figure 2C**). Therefore, we move on to the functional test of EV-siRNA delivery.

**Figure 2.**
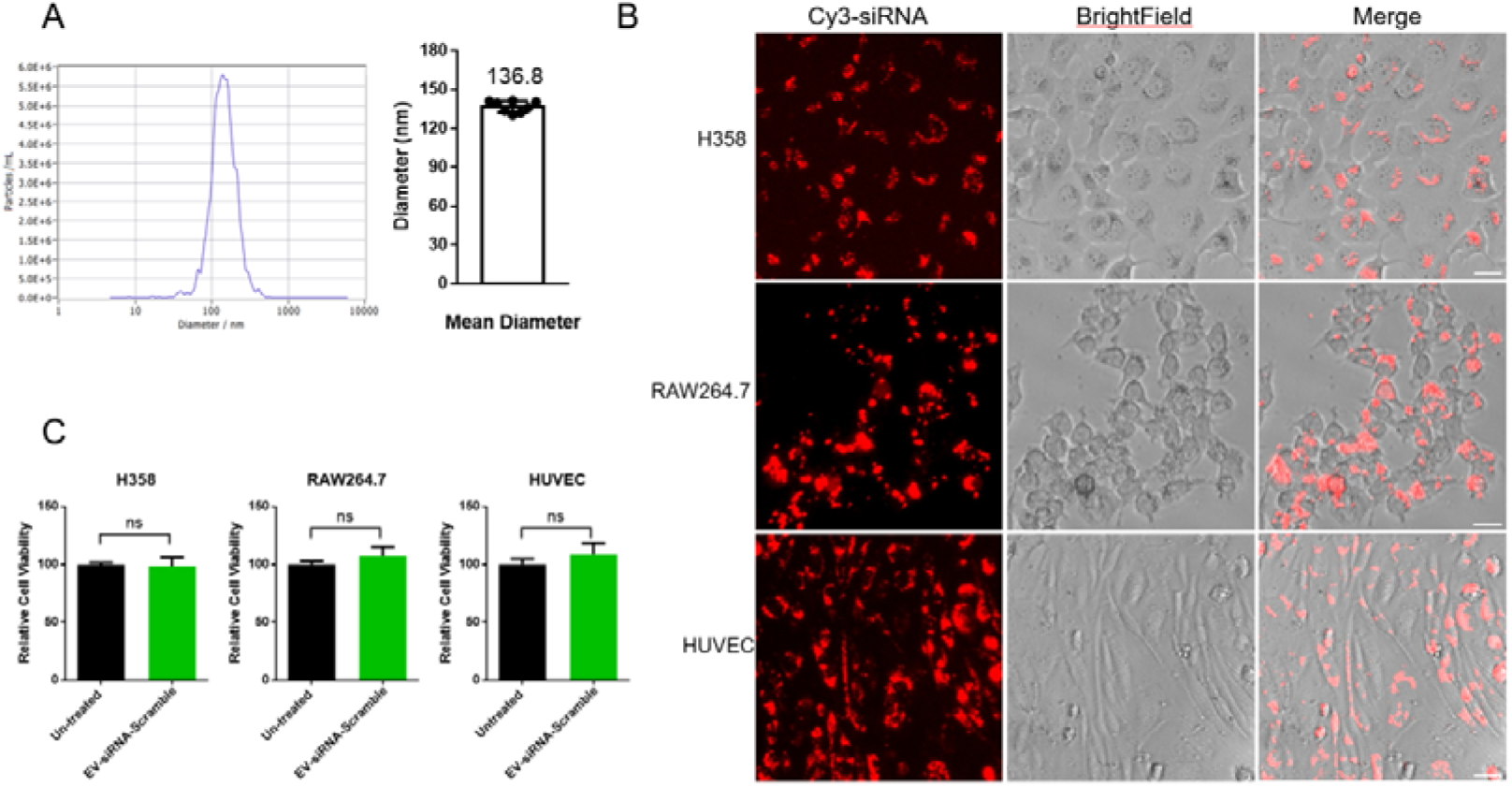
Characterization of extracellular vesicles loaded with siRNA. (A) Representative nanoparticle tracking analysis (NTA) analysis of size distribution and concentration of EVs loaded with siRNA. The mean diameter from eight different batches of EV-siR were presented. (B) Representative images of cellular uptake of cy3 labeled EV-siR (final siRNA concentration: 50 nM). Scale bar: 5 μm. (C) Cell viability analysis of cells at 72 hours after transfection with EV-siR-Scramble (final siRNA concentration: 50 nM). N=3. ns: not significant.

### In vitro EV-siRNA delivery

To examine if the observed EV-siRNA uptake in cells can lead to efficient gene silencing, siRNA for tumor oncogene *KRAS* was designed and efficient knockdown was observed at 48 hours post transfection (**Figure 3A**). We further selected a verified model siRNA siR-PT, in which the target gene was essential for cell survival. As expected, EV-siR-PT transfection resulted in significant cellular toxicity in several cells tested, as shown by both microscopy examination and CCK-8 assay (**Figure 3B-C**). Next, to evaluate the stability of EV-siRNA, we tested gene knock down efficiency of EV-siRNA after short and long-time storage. Remarkably, EV-siRNA retained similar *KRAS* knock down efficiency after storage at 4 degrees for one week and even for four weeks (**Figure 3D**). Overall, EV-siRNA demonstrate efficient delivery *in vitro*.

**Figure 3.**
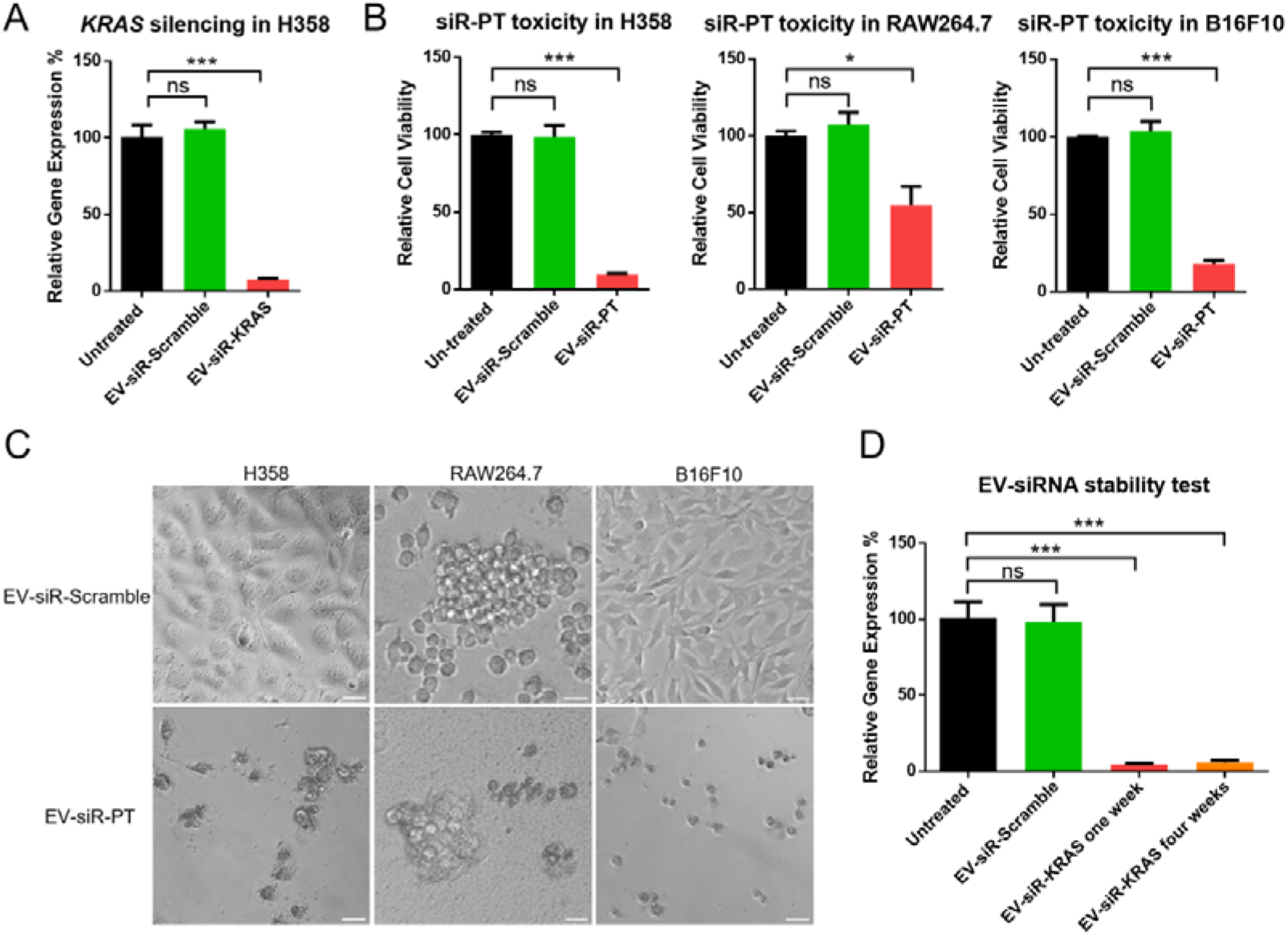
*In vitro* EV-siRNA delivery analysis. (A) RT-qPCR analysis of relative *KRAS* knock down efficiency at 48 hours post EV-siR transfection (final siRNA concentration: 50 nM). N=3. (B) Cell viability analysis at 48 hours post EV-siR transfection (final siRNA concentration: 10 nM). N=3. (C) Representative microscopy images of cells at 48 hours post EV-siR transfection (final siRNA concentration: 10 nM). Scale bar: 5 μm. (D) RT-qPCR analysis of *KRAS* knock down efficiency at 48 hours post EV-siR-KRAS transfection (final siRNA concentration: 50 nM). EV-siRNA were stored at 4 degrees for one week or four weeks. N=3.ns: not significant; *: p<0.05; ***: p<0.001.

### In vivo EV-siRNA delivery

Since EV-siRNA demonstrate excellent efficiency *in vitro*, we next set out to evaluate if EV-siRNA are able to function consistently *in vivo*. B16F10 melanoma xenograft model was established in C57BL6/N mice background, and EV-siR-PT were intratumorally administrated to see if EV-siR-PT could deliver the toxic siRNA and inhibit tumor growth (**Figure 4A**). In the control group, as expected, the B16F10 tumor xenograft expanded rapidly, recorded more than 35-fold expansion in one week as well as in parallel treated EV-siR-Scramble groups (**Figure 4B**). In contrast, EV-siR-PT significantly inhibited tumor growth, which achieved about 50% tumor inhibition as measured by tumor volume and tumor mass (**Figure 4B-D**). Therefore, we show in this study that EVs are able to deliver siRNA efficiently *in vivo* as well.

**Figure 4.**
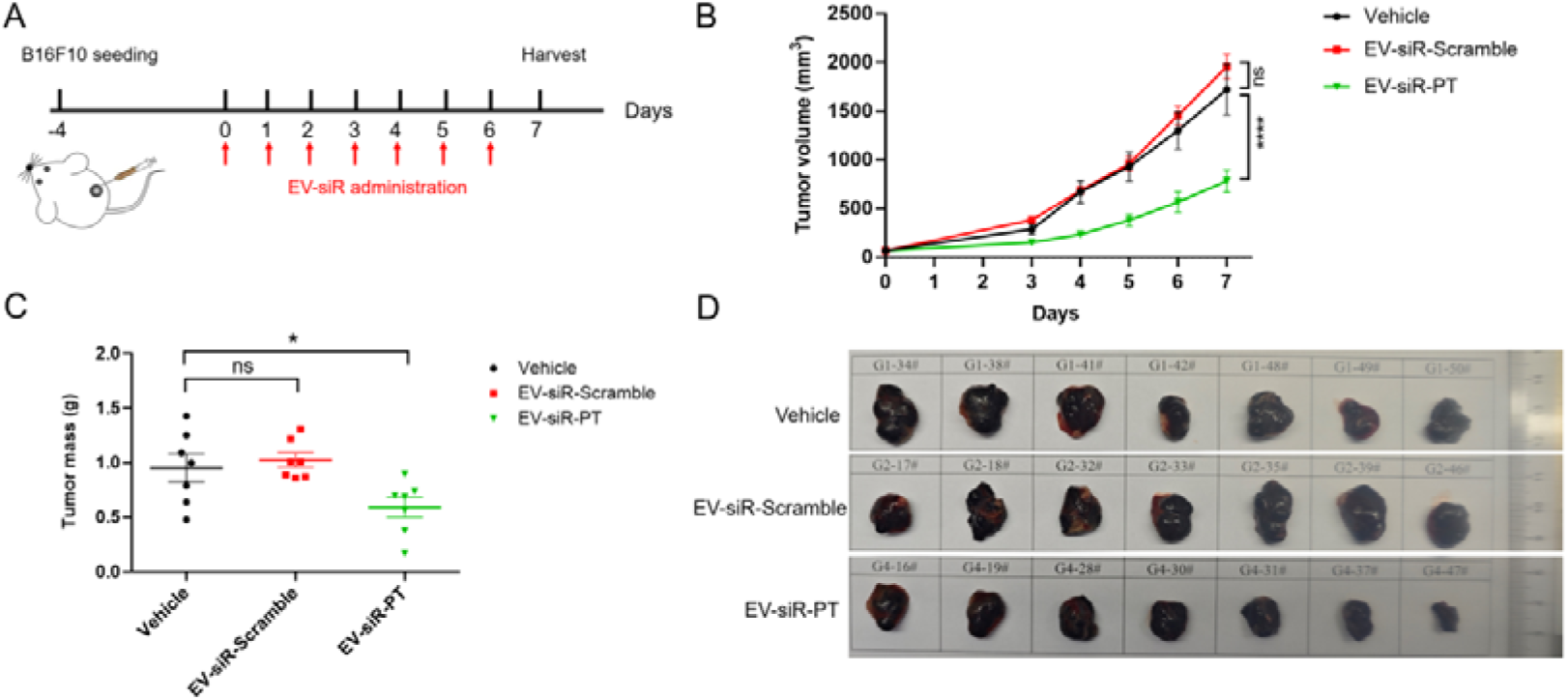
*In vivo* EV-siRNA delivery analysis. (A) Schematic of B16F10 tumor xenograft model in C57BL6/N mice. (B) Tumor volume tracking for mice receiving different treatment groups. N=7. (C) Tumor mass measurement at end point for different treatment groups. N=7. (D) Images of tumor at end point for different treatment groups. ns: not significant; *: p<0.05; ****: p<0.0001.

## Discussion

Extracellular vesicles have been explored as potential siRNA delivery platform in several studies, yet further development is still hampered by low quality preparation and inefficient siRNA loading (Alvarez-Erviti et al. 2011; El-Andaloussi et al. 2012; Kumar et al. 2015; Kamerkar et al. 2017). In this work, we have provided solutions to overcome current limitations. Firstly, through combining high density HEK293F cell culture and density gradient based EVs purification method, sufficient EVs are produced in high quality. To date, this is the cleanest EVs preparation ever presented. An innovative method other than electroporation is further developed to load siRNA into EVs, and functional testing in both *in vitro* and *in vivo* models have validated the excellent efficiency. In summary, we have established efficient EVs platform for siRNA delivery.

Currently, GalNAc-siRNA conjugates is the dominant platform for siRNA therapeutics, yet it’s only applicable for liver associated diseases. An siRNA delivery platform with extra-hepatic targeting ability is urgently needed to fully unleash the power of siRNA therapeutics, especially for those valuable, undruggable targets by small molecule or antibody drugs. In this study, we have shown the successful and efficient delivery of EV-siRNA to a variety of cell types including primary cells and tumor cells. Furthermore, EV-siRNA delivered to tumor xenograft model significantly inhibit tumor growth. In addition, previous studies have demonstrated that EVs can further target a specific cellular population when surface modified with targeting peptides or antibodies (Zeng et al. 2023). The unique properties of exosomes make them a promising avenue for advancing siRNA delivery technology in a safe and effective manner.

## Materials and Methods

### Cell culture

HEK293F cells were purchased from Quacell. H358, B16F10, RAW264.7 and HUVEC cells were purchased from Procell. HEK293F cells were cultured in OPM-CD05 medium (OPM Biosciences). H358 and B16F10 cells were cultured in RPMI1640 medium (Gibco) supplemented with 10% feta bovine serum (Gibco) and 1 × penicillin-streptomycin (Gibco). HUVEC cells were cultured in DMEM medium (Gibco) supplemented with 10% feta bovine serum and 1 × penicillin-streptomycin. Suspension cell culture were maintained on orbital shaker (90 RPM) in a humidified incubator at 37□ with 8% CO2. Adherent cell culture were maintained in a humidified incubator at 37□ with 5% CO2.

### Purification of extracellular vesicles (EVs)

The cell culture supernatant was first centrifuged at 133,900 g at 4 □ for 60 min. The crude EVs pellet were resuspended in PBS, and further layered onto 17.5% iodixanol/45% iodixanol gradient, followed by centrifugation at 150,000 g at 4 □ for 16 h. The extracellular vesicles appeared as a white layer between PBS/17.5% iodixanol, which were carefully pipetted out and washed with PBS by centrifugation at 135,000 g at 4 □ for 3 h. The refined EVs were finally resuspended in PBS.

### Western blot antibodies

The following antibodies were used: CD9(PTG, 60232-1-Ig), CD63(PTG, 25682-1-AP), CD81(PTG, 66866-1-Ig), TSG101(BD, 612696), beta Actin(ThermoFisher, MA5-16410), HRP Goat Anti-Rabbit IgG (H+L) (ABclonal, AS014), HRP Goat Anti-Mouse IgG (H+L) (ABclonal, AS003).

### Transmission Electron Microscopy

Extracellular vesicles were applied onto the glow-discharged copper grid (200 mesh, coated with carbon film). To perform negative staining, 2% uranyl acetate were incubated with EVs at room temperature for 1 min, followed by quick wash with distilled water to remove excess stain. The grids were air-dried before being imaged under Tecnai G2 transmission electron microscope (Thermo FEI, 120 kV).

### Nanoparticle tracking analysis (NTA)

To evaluate the size distribution and concentration of extracellular vesicles, EVs in PBS solution were applied directly into ZetaView Nano Particle Tracking Analyzer (ParticleMetrix, PMX120-Z).

### RT-qPCR

All siRNA were synthesized by Sangon Biotech. Forty-eight hours post EV-siRNA treatment, total RNA from the cells plated on 96 well plates were first extracted and purified using the RNAprep Pure Micro Kit (TIANGEN). Reverse transcription was performed with HiScript III cDNA Synthesis Kit (Vazyme). Next, the cDNA was added into the 2 × SYBR green qPCR mix (Ezbioscience), and quantitative PCR were analyzed with QuantStudio3 (Applied Biosystems).

### CCK-8 assay

To perform CCK-8 assay, 10 μL of CCK-8 reagent (Vazyme) were added to the cells directly in 96 well plates. After incubation at 37 □ incubator for two hours, absorbance at 450 nm was recorded on microplate reader (Pherastar).

### Tumor xenograft model

Female, 8-weeks-old C57BL6/N mice (Vital River) were implanted with 1E6 B16F10 cells/mice under the right fat pad region. When the average tumor volume reached around 60 mm^3^, the mice were randomly grouped for different treatment conditions. Intratumoral injections and tumor volume measurement were performed every day for seven days consecutively. For first three days, mice were injected with 25 μL PBS or 50 nM EV-siRNA in PBS; for the last four days, mice were injected with 50 μL PBS or 50 nM EV-siRNA in PBS. On last day, mice were sacrificed and tumors were excised out and imaged.

## Notes

### Competing Interest Statement

All authors have interests in Vesicure Therapeutics Co. Ltd.

